# Dopamine increases accuracy and lengthens deliberation time in explicit motor skill learning

**DOI:** 10.1101/2023.01.31.526542

**Authors:** Li-Ann Leow, Lena Bernheine, Timothy J Carroll, Paul E Dux, Hannah L Filmer

## Abstract

Although animal research implicates a central role for dopamine in motor skill learning, a direct causal link has yet to be established in neurotypical humans. Here, we tested if a pharmacological manipulation of dopamine alters motor learning, using a paradigm which engaged explicit, goal-directed strategies. Participants (27 females, 11 males, aged 18-29 years) first consumed either 100mg of Levodopa (n=19), a dopamine precursor that increases dopamine availability, or placebo (n=19). Then, during training, participants learnt the explicit strategy of aiming away from presented targets by instructed angles of varying sizes. Targets shifted mid-movement by the instructed aiming angle. Task success was thus contingent upon aiming accuracy. The effect of the dopamine manipulations on skill learning was assessed during training, and at an overnight follow-up. Increasing dopamine availability improved aiming accuracy and lengthened reaction times, particularly for larger, more difficult aiming angles, both at training, and at follow-up. Results support the proposal that dopamine is important in decisions to engage instrumental motivation to optimise performance, particularly when learning to execute goal-directed strategies in motor skill learning.

---

Motor skill learning is essential for survival: a hungry bear catching salmon must adapt its paw-strikes to dynamic task parameters, such as the movement and friction of the salmon’s body and the forces applied by river waters. Reward is a potent modulator of skilled movement, affecting movement in two primary ways. First, rewards can increase the vigor of movements (e.g.,Manohar et al., 2015; Carroll et al., 2019a; Codol et al., 2019), even when rewards are not performance-contingent (Takikawa et al., 2002). Second, rewards can boost skill learning. For example, rewarding participants for achieving performance criteria can speed-up learning (Vassiliadis et al., 2021) or improve retention of motor learning (Abe et al., 2011; Madelain et al., 2011; Galea et al., 2015). The potency of reward in altering behaviour and learning is perhaps unsurprising, as animals must learn motor skills to attain resources for survival (Barron et al., 2010).

Experimental manipulations of reward do not, however, always benefit motor learning (e.g., Steel et al., 2016; Spampinato et al., 2019). This is perhaps because rewarding participants for achieving some experimenter-determined performance criterion does not directly alter a key learning signal: success or failure at achieving the task goal. Failures to achieve task goals, termed task errors, seem essential to forming memories that improve future motor performance, and such memories take at least two forms: deliberative goal-oriented strategies, and automatic stimulus-response associations (Leow et al., 2020).

Task errors can be conceptualised as a form of reward prediction error (Leow et al., 2020), which is a broader term describing discrepancies between predicted rewards and received rewards (Sutton and Barto, 1998). Transient bursts of firing by midbrain dopamine neurons trigered by reward prediction errors have been thought to “stamp-in” associations between a stimulus and its associated response, resulting in the formation of automatic stimulus-response mappings. Increasing evidence also implicates dopamine in the engagement of deliberative, goal-oriented behaviours (e.g., Akam and Walton, 2021). Dopamine might therefore play dual roles in skill learning: stamping-in stimulus-response associations to facilitate automaticity with training, and motivating the engagement of goal-oriented strategies.

Whilst non-human animal studies implicate dopamine-dependent circuits in skill learning (e.g., Hosp et al., 2011), inter-species differences in the cognitive and neural processes associated with the dopamine system (e.g.,Khan et al., 1998) limits generalisation of findings from animals to humans. Evidence for the role of dopamine in human motor learning comes from studying how dopamine medications alter learning in patients with impaired dopamine function (e.g., Kwak et al., 2010), but heterogeneity in disease phenotypes complicates inferences from such work (Cools et al., 2022). Causal evidence for the role of dopamine in skill learning in neurotypical humans is scarce, and limited to studies of elementary motor tasks such as repeating simple thumb-abduction movements (Floel et al., 2008; Rösser et al., 2008), or tracking on-screen targets (Chen et al., 2020a). Of import, some studies have found no effect of manipulating dopamine on motor learning (Quattrocchi et al., 2018; Palidis et al., 2021). These null results might have resulted from paradigms which predominantly engage implicit learning (Palidis et al., 2021), or which do not dissociate effects of explicit and implicit learning proceseses (Quattrocchi et al., 2018), which can have mutually compensatory effects on behaviour (Albert et al., 2022). Indeed, in reinforcement learning tasks, explicit, goal-directed processes can be more sensitive to effects of dopamine manipulations than implicit, automatic processes (Sharp et al., 2016). Similarly, explicit processes are more sensitive to reward manipulations than implicit processes in motor learning (Holland and Codol, 2017; Codol et al., 2018). Manipulations of dopamine and reward might thus be more observable in learning tasks driven by explicit knowledge of the task structure and strategies to achieve task goals.

Here, we investigated the role of dopamine in learning goal-directed motor strategies. After consuming the dopamine precursor Levodopa or placebo, participants learnt a strategy of aiming away from presented targets by angles of varying sizes (e.g.,Georgopoulos and Massey, 1987a). Participants had to employ strategic aiming to hit targets, as targets would jump mid-movement by the instructed aiming angle. Thus, task success (i.e., target-hitting) was contingent upon successful aiming. Effects of the dopamine manipulations were assessed during training and at a no-drug follow-up session.

## Method

### Participants

Thirty-eight neurotypical young adults (27 females; median age = 21.58 years; range= 18-29 years) were recruited from The University of Queensland community, and were reimbursed for participation (AUD$20/hour). Participants were screened for neurological and psychiatric conditions and contraindications for Levodopa, and provided written informed consent. In accordance with the National Health and Medical Research Council’s guidelines, this experiment was approved by the human ethics committee at The University of Queensland. No datasets were excluded from the study.

As no published studies on the effects of Levodopa on explicit motor skill learning in neurotypical adults existed at the time of planning the study, we based our sample size on previous studies that examined effects of Levodopa on tasks requiring cognitive effort (Vo et al., 2017). Half of the participants (n=19) were randomly assigned to the Levodopa condition (16 females; median age = 20.74 years; range = 18-25 years), the other half were assigned to the Placebo condition (n=19, 11 females; median age = 22.42 years; range = 19-29 years).

### Drug manipulation

The Levodopa group consumed the dopamine precursor, a Madopar® 125 (Levodopa 100mg/ Benserazide 25mg) tablet, whereas the Placebo Group consumed a crushed multivitamin (Centrum® for women). Double blinding was achieved by having an experimenter who was not involved in data collection crush and disperse the tablets in orange juice before consumption by the participant. Behavioural testing of the aiming task began approximately 60 minutes after tablet administration, around the time of peak plasma concentration (Contin and Martinelli, 2010). The drug manipulation was only employed on the training session, and not at follow-up.

### Apparatus

A vBot planar robotic manipulandum was used (Howard et al., 2009). This apparatus has a low-mass, two-link carbon fiber arm and measures position with optical encoders sampled at 1,000 Hz. Participants completed the aiming task while seated on a height-adjustable chair at their ideal height for viewing the screen. Visual feedback was presented on a horizontal plane on a 27” LCD computer monitor (ASUS, VG278H, set at 60-Hz refresh rate) mounted above the vBOT and projected to the subject via a mirror in a darkened room. Participants had no direct vision of their hand. Hand position was represented by a red cursor (0.25 cm radius circle) on a black background. The mirror also allowed visual feedback of the target (a 0.5 cm-radius circle) and the start location (a 0.35 cm radius circle), which was aligned 10 cm to the right of the participant’s midsagittal plane at approximately mid-sternum level.

### Procedure

Participants were first given the following instructions: “You are about to perform sets of 80 trials in which you will be asked to move at specific angle relative to the target in clockwise direction. The angle will differ between blocks. We will tell you what the angle is for each block. First, make up your mind and then try to keep your movement straight and fast. Try your best to execute your movements as accurately as possible and hit as many targets as possible. Between each set we will ask you to aim towards the target in a 0° angle, that means you should hit the target directly. These sets will contain 20 trials.” Instructions were followed by a demonstration by the experimenter on how to re-aim by 90° away from the presented target.

Each trial commenced by displaying the central start circle, with participants then moving the cursor within 1 cm of the start circle. If participants did not move to the start after 1s, the robotic manipulandum moved the participant’s hand to the start circle, using a simulated 2-dimensional spring with the spring constant magnitude linearly increasing over time. After the cursor remained within the start circle at a speed below 0.1 cm/s for 200 ms, targets appeared in random order at one of eight locations 9 cm away from the start circle (0°, 45°, 90° … 315°) (see Figure 1A).

**Figure 1A:**
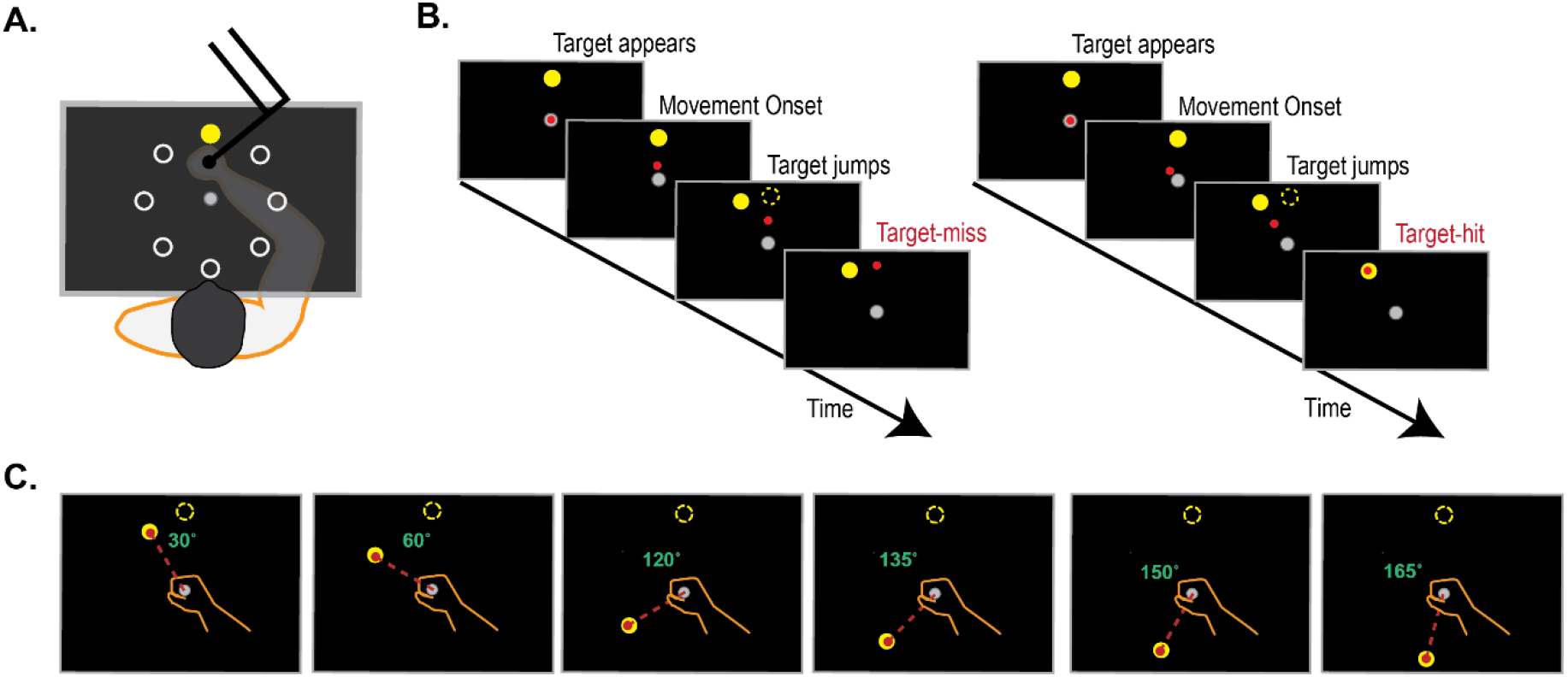
Participants used a robotic manipulandum to move an on-screen cursor from a central start circle to one of 8 possible target locations (0°, 45°…315°): target order was randomised. B. Trial sequence for strategic aiming trials. In blocks of 80 trials, participants were instructed to aim away from the presented target at angles of varying sizes (0°, 30°, 60°, 120°, 135°, 150°, 165°, see C). Strategic aiming was required to successfully hit the target, as targets jumped mid-movement (4cm into the 9cm start-target distance) by the instructed angle.

Participants were given seven aiming blocks (1 block = 80 trials), where they were asked to re-aim away from the presented target by one of the following angles: 0°, 30°, 60°, 120°, 135°, 150°, 165°, similar to previous work (Georgopoulos and Massey, 1987b; Pellizzer and Georgopoulos, 1993; Bhat and Sanes, 1998; Neely and Heath, 2010; McNeely and Earhart, 2012). Block order was randomised between subjects. During each aiming block, the target “jumped” mid-movement (i.e., when movement extent reached 4cm into the 9cm start-target distance): the size of the target-jump was the same as the instructed aiming angle, such that successful aiming was necessary to attain the task goal of hitting the target (Leow et al., 2020). Target “jumps” were achieved by extinguishing the target and re-displaying the target at the new location as soon as movement extent exceeded 4cm out of the 9cm start-target distance (see Figure 1B). Each aiming block was followed by a 20-trial washout block, where the task was to move straight to the presented targets. To economise study duration, a washout block was not provided after the 0° block or the final re-aiming block. Across all trials, cursor feedback was only given after movement extent exceeded 4cm, and was then extinguished after movement extent exceeded the 9cm start-target distance. Across all trials, including the washout blocks, auditory feedback in the form of a beep sound (coin.wav from Super Mario) was given when the reach angle measured at 4cm into the 9cm start-target distance reached an accuracy criterion of being within a +/10° range of the ideal aiming angle. This auditory feedback was implemented to maintain participant engagement in the long test session.

After an overnight delay, participants returned for a follow-up session, scheduled a minimum of 18 hours after the first session. The task was identical to the first session. On the second session, all participants consumed a placebo pill dispersed in orange juice. Participant blinding was assessed after both sessions.

### Additional assessments

To assess for changes in arousal, heart rate, blood pressure, and mood these variables were measured at the beginning, after one hour and at the end of each session (Vo et al., 2016). Mood was assessed using the Mood Rating Scale (Bond and Lader, 1974) which includes sixteen items separated in following factors: alertness, contentedness and calmness. The participants could rate each element within a range of 10 points. The three factors were evaluated as a total score.

Individual differences in baseline dopamine function affects responsivity to dopamine medications. Individual dopamine baseline function is predicted by impulsivity and working memory. Impulsivity, as measured by the behavioural inhibition scale (BIS-11) is associated with dopamine D2/D3 receptor availability in the midbrain (Buckholtz et al., 2010). BIS scores can predict effects of methylphenidate on learning (Clatworthy et al., 2009). Similarly, working memory span has been associated with dopamine synthesis capacity, with a medium to large effect size for the correlation between listening span scores and dopamine synthesis capacity (Cools et al., 2008). Working memory span can predict effects of pharmacological manipulations of dopamine (Broadway et al., 2018; Fallon et al., 2019). To account for individual differences in baseline dopamine function, scores on assessments of impulsivity and working memory capacity were used as covariates. Working memory was measured with the memory updating task from a validated working memory battery (Lewandowsky et al., 2010). The memory updating task had participants remember a set of digits presented for 1 second in separate on-screen locations, and update these digits through arithmetic operations shown on-screen for 1.3 seconds at corresponding locations. The number of digits presented (i.e., set size) varied from three to five across trials. After a varying number of update operations (between two and six), question mark prompts appeared in each frame. There was no time limit for the recall, and there was no performance feedback. The operations ranged from +7 to −7, excluding 0, and the results from 1 to 9. There were 15 trials in total and 2 practice trials. Performance on the memory updating task was assessed both at training and at follow-up, but to avoid the influence of practice effects, memory updating data from only the training session were used for analysis.

### Data analysis

Trials were first grouped into bins of 8 (1 visit to each of the 8 targets per bin) for analysis. To evaluate how overall performance and improvements across session-to-session were affected by Levodopa and by the varying levels of aiming difficulty, we ran analyses on the following dependent variables: (1) **Reach accuracy,** defined as the difference between the ideal aiming direction and the absolute values of the reach direction, measured at 20% of the start-target distance (i.e., 1.8 cm into the 9cm start-target distance). Note that for the 0° blocks, this estimation of accuracy resulted in negative values for reaches that deviated from the straight ahead direction. (2) **Reaction time**, defined as the interval from target appearance to movement onset. Movement onset was defined as the time at which the hand speed first exceeded 2 cm/s. (3) **Reach variability**, defined as the standard deviation of reach directions every bin (every 8 trials). (4) **Reaches meeting criteria for auditory feedback,** defined as the proportion of reaches per bin which were within 10 degrees of the ideal aiming angle at 4cm into the 9cm movement. In addition to the primary task success manipulations, we provided auditory feedback (Mario coin.wav) to maintain participant engagement when reaches met the performance criteria. This auditory feedback might be interpreted as a form of extrinsic reward, and thus we additionally explored of the dopamine manipulations altered the number of reaches meeting criteria for auditory feedback.

To quantify whether the drug manipulation altered learning and performance of the task, Bayesian analyses of variances with the within-subjects factors Session (Session 1, Session 2) x Aiming Angle (0°, 30°, 60°, 120°, 135°, 150°, 160°) and Bin (1, 2,…10) and the between-subjects factor Drug (Levodopa, Placebo) were run on variables of interest. Working memory capacity (estimated via memory updating scores), and impulsivity (estimated via BIS total scores) were entered as covariates. As gender affects performance in mental rotation tasks (Terlecki et al., 2008), which share processes with the visuomotor mental rotation task used here, we also entered gender as a covariate of no interest. We used Bayesian statistics to quantify evidence for the test hypothesis, as well as evidence for the null hypothesis. Inclusion Bayes factors (BFinclusion, BF_incl_) were determined to estimate the strength of evidence in favour of including an effect relative to models stripped of that effect. Exclusion Bayes Factors (BFexclusion, BF_excl_) were determined to estimate the strength of evidence in favour of excluding an effect relative to models stripped of that effect. In post-hoc tests, where evidence for the null hypothesis is reported as BF_01_, and evidence for the test hypothesis is reported as BF_10_, the posterior odds were corrected for multiple testing by fixing to 0.5 the prior probability that the null hypothesis holds across all comparisons (Westfall, Johnson, & Utts, 1997). Jeffreys’s evidence categories for interpretation (Wetzels et al., 2011), were taken as the standard for evaluation of the reported Bayes factors. Specifically, Bayes factors of 1–3 were interpreted as anecdotal, 3–10 as moderate, and greater than 10 as strong evidence for the test hypothesis, whereas Bayes Factors of 0.33 or less was taken as evidence for the null hypothesis. Where appropriate, cohen’s d values were used to quantify effect sizes, and were reported with 95% confidence intervals of the effect size.

## Results

### Control measurements

Pharmacological manipulations of dopamine can elicit undesired side effects such as nausea, resulting in some participants withdrawing from the study (for example, see Chen et al., 2020a), and can also change mood state (Vo et al., 2018). None of our participants reported nausea, and all participants completed the study. Levodopa did not alter our participants blood pressure or mood, as Time (before drug administration, 1 hour after drug administration, 2 hours after drug administration) x Drug ANOVAs yielded evidence for the null hypothesis for Time x Drug interactions for blood pressure (diastole: BF_excl_ = 6.383, systole: BF_excl_ = 5.683) and mood (BF_excl_ = 3.681). Heart rate was reduced to a greater extent for participants on placebo (mean reduction in heart rate: 10.211+/- 1.828) than levodopa (mean reduction in heart rate: 1.737+/- 1.828), as shown by a Time x Drug interaction, BF_incl_ = 6.519).

Participants were not above chance at accurately guessing the drug condition, as shown by Bayesian binomial tests (BF0+ = 4.685).

### Manipulation check

Performance scaled with task difficulty, as larger aiming angles decreased reach accuracy, increased variability and lengthened reaction times (see Figure 2), as supported by main effect of Angle for accuracy (BF_incl_ = 2.25E+117), error variability (BF_incl_ = ∞) and reaction times (BF_incl_ = ∞), and trials meeting criteria for auditory feedback (BF_incl_ = ∞). In contrast to motor learning paradigms involving both implicit and explicit learning processes, which show trial-by-trial improvements in performance (either in increased accuracy and/or decreased reaction times), here, performance was similar across trial bins for accuracy and reaction time. Only reach variability showed evidence for a main effect of bin (BF_incl_ = 66600), where reach variability was large in the first bin, but rapidly reduced from the first to the second bin (post-hoc t-tests comparing the first to the second bin, BF_10_ = 31.945), but did not reduce after the second bin (all BF_01_ greater than 5). This large variability in the first bin, followed by a reduction in variability in all subsequent bins, and the absence of gradual performance improvements across bins for accuracy or reaction times, is consistent with previous characterisations of explicit motor learning (e.g., Benson et al., 2011).

**Figure 2.**
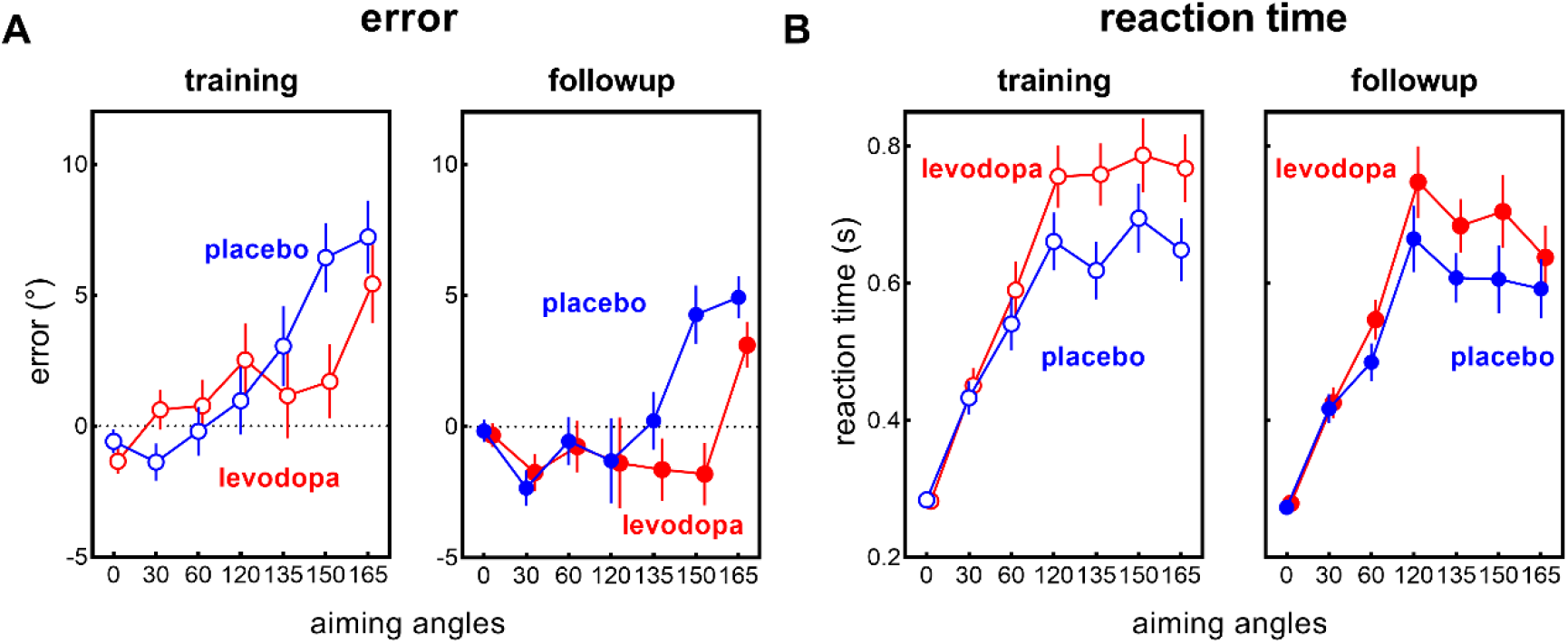
At larger aiming angles, errors were smaller with Levodopa than with placebo (A), and reaction times were longer with Levodopa than with placebo (B), both at training (clear symbols) and follow-up (filled symbols). Values are estimated marginal means and standard errors of the mean.

**Figure 3.**
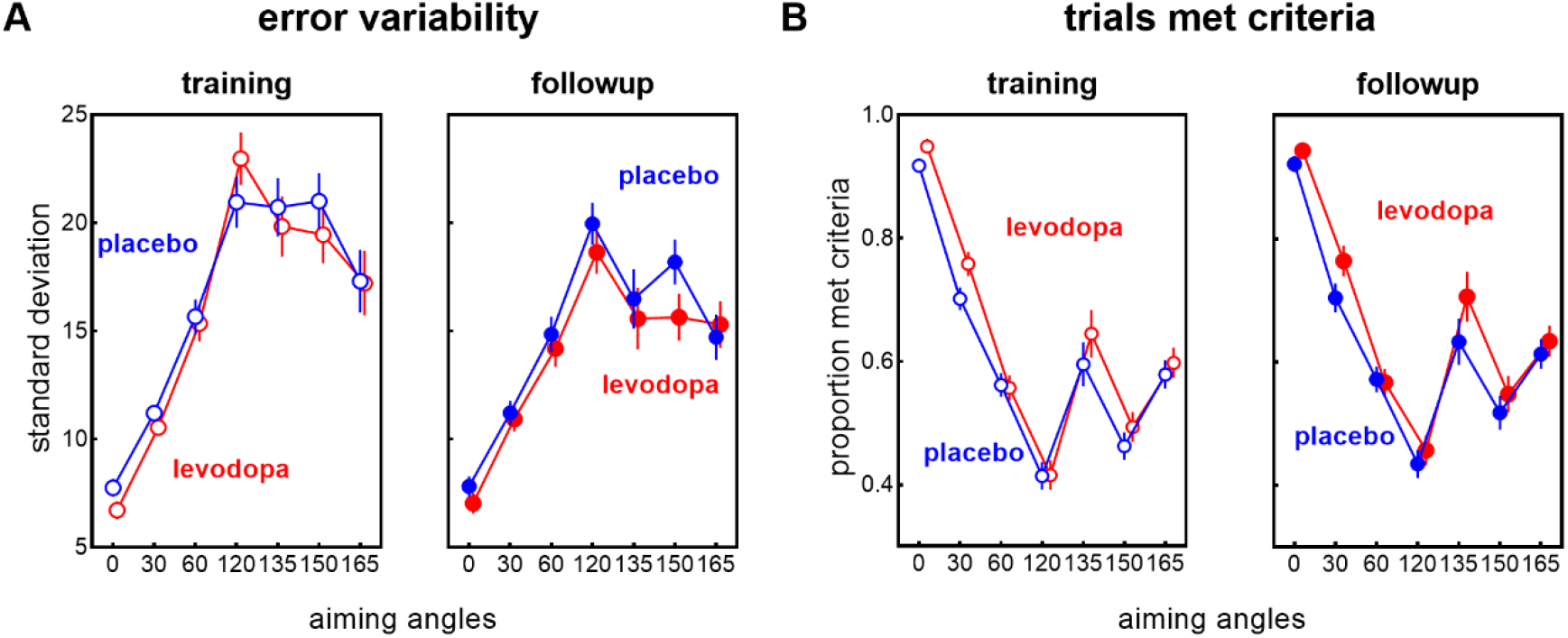
Levodopa did not alter error variability (A) or proportion of reaches meeting criteria for auditory feedback (B), both at training and follow-up. Values are estimated marginal means and standard errors of the mean.

### Levodopa increased accuracy whilst lengthening reaction times under difficult conditions

Levodopa administered before training altered performance for difficult aiming angles, at both training and follow-up (see Figure 2), as shown by Angle x Drug interactions for accuracy [BF_incl_ = 3.89E+12] and reaction times [BF_incl_ =2.61E+11]. Levodopa increased accuracy, and this effect was most prominent with the larger aiming angles, see Figure 2A. Analyses comparing the Levodopa and placebo groups for different rotation angles showed comparatively larger effect sizes for larger angles, such as for 150° (cohen’s d = −0.628, 95% CI [−1.284, 0.027]) and 165° (cohen’s d = −0.158, 95% CI [−0.759, 0.442]) than for smaller angles such as 30° (cohen’s d = 0.165, 95%CI [−0.436, 0.766]) and 60° (cohen’s d = 0.065 [−0.532, 0.663])

Levodopa also lengthened reaction times, particularly with larger aiming angles, see Figure 2B. Analyses comparing the Levodopa and placebo groups for different rotation angles showed larger effect sizes for larger angles, such as for 150° (cohen’s d = 0.58, 95% CI [−0.609,1.769]) and 165°(cohen’s d = 0.499, 95% CI [−0.681,1.679]) than for smaller angles, such as 30° (cohen’s d = 0.08, 95% CI [−1.074,1.235]) and 60° (cohen’s d = 0.336, 95% C [−0.829,1.502]).

Thus, with larger aiming angles, as the task became more difficult, the impact of Levodopa increased: increasing accuracy whilst lengthening reaction times. In a task where success depended on aiming accuracy, increasing dopamine availability via Levodopa resulted in a tendency to trade speed for accuracy.

### No effect of Levodopa on reach variability and proportion of trials meeting criteria for auditory feedback

As reward can alter movement precision (Manohar et al., 2015), we examined if our dopamine manipulations altered movement precision, quantified via reach variability. Levodopa did not alter reach variability, as the main effects of drug (BF_excl_ = 4.642), the drug x angle interaction (BF_excl_ =190.585) and the session x drug x angle interaction (BF_excl_ = 175.627) showed evidence for the null hypothesis.

Previous null findings with dopamine drug manipulations on motor learning gave participants extrinsic rewards contingent upon successful performance (Quattrocchi et al., 2018; Palidis et al., 2021). In addition to our target manipulations which putatively acted upon rewards associated with achieving the task goal (target-hitting), we also gave participants auditory feedback when a performance criterion was met (reach errors < 10°). This feedback might be interpreted by participants as a form of extrinsic auditory reward. Levodopa did not increase the proportion of trials which met the criterion for auditory feedback, as there was evidence to support absence of the main effect of drug (BF_excl_ = 5.174) and absence of a drug by angle interaction (BF_excl_ =11.763). Thus, whilst Levodopa resulted in an increase in accuracy for larger aiming angles, this increase in accuracy was less than 10° (see Figure 2), and Levodopa did not increase the number of trials that met the criterion for potentially rewarding auditory feedback.

### Levodopa did not alter session-to-session performance improvements

Performance improvements from the training session to the follow-up session were evidenced in greater accuracy (lower error), faster reaction times, lower error variability, and more trials meeting the criterion for auditory feedback (see Figure 4). This was supported by clear evidence for the main effect of session on accuracy (BF_incl_ = 2.90e+16), reaction times (BF_incl_ = 2.21E+35), precision (BF_incl_=3.51e+16), and trials achieving criterion for auditory feedback (BF_incl_ = 5.591e+14). Levodopa did not, however augment offline performance improvements, as shown by evidence for the null hypothesis for Session x Angle x Drug interactions (accuracy: BF_excl_ = 2548.451, error variability: BF_excl_ = 175.627, trials meeting criterion for auditory feedback, BF_excl_ = 351.036). For reaction time, there was moderate evidence for a Session x Angle x Drug interaction, BF_incl_ = 4.065, where the reduction in reaction time from training to follow-up for the two largest angles tended to be larger for the levodopa group (mean difference = 90ms [11ms, 170ms] than the placebo group (mean difference = 53ms [−8ms, 147ms]).

**Figure 4.**
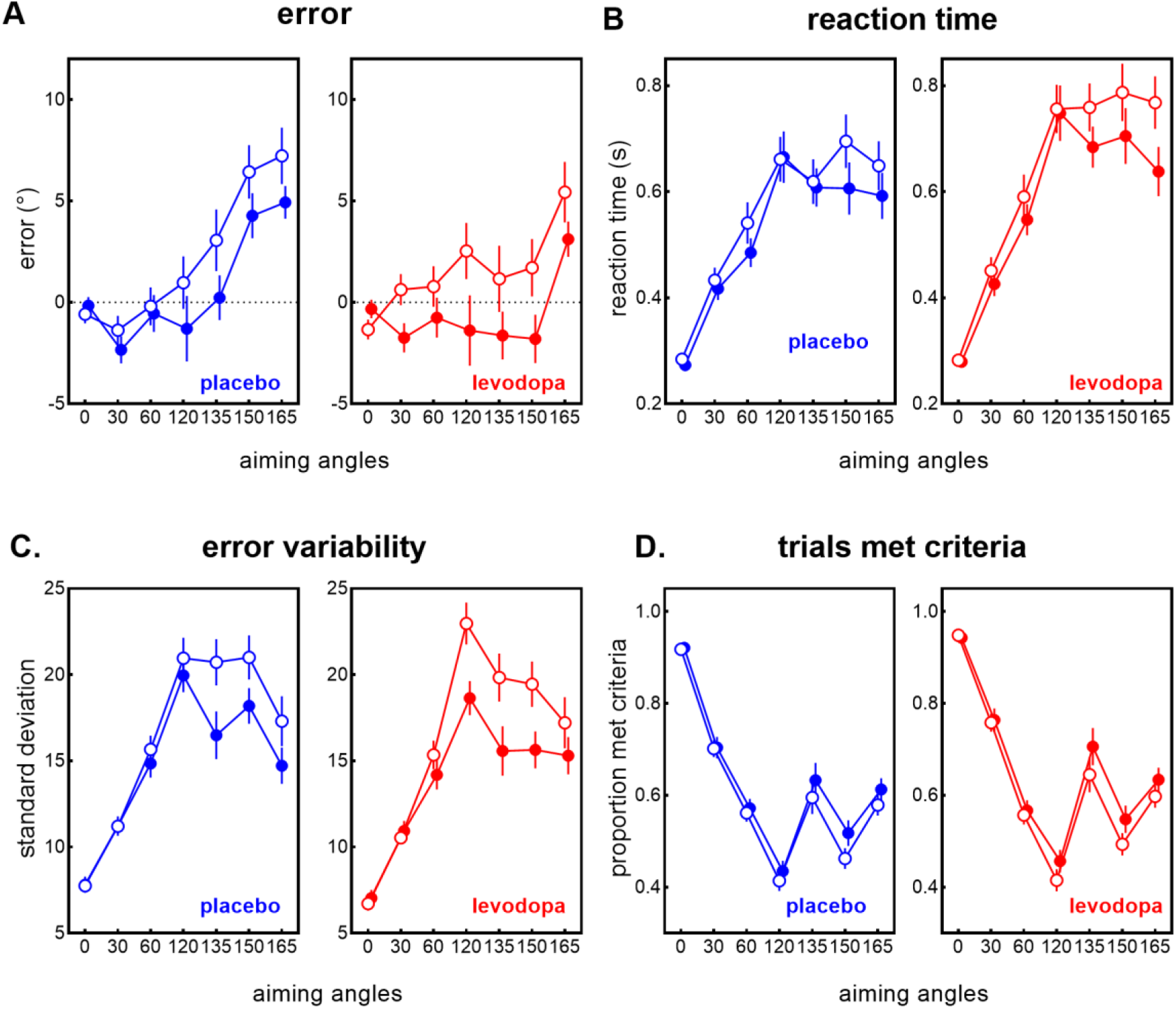
Performance improvements at follow-up (filled circles) compared to training (open circles), as evidenced in increased accuracy (decreased errors) (A), shorter reaction times (B), reduced error variability, and more trials meeting criteria for auditory feedback (D). Values are estimated marginal means and standard errors of the means.

### Effect of task difficulty and levodopa on movement vigor

The relationship between cognitive and movement vigor has been underexplored (for a recent review, see Shadmehr et al., 2019). As part of a secondary, exploratory analysis, we tested how completing a task of varying cognitive difficulty within the context of an explicit motor learning task affected movement vigor, and whether increasing dopamine availability alters the way task difficulty affects movement vigor. We quantified vigor using (1) peak velocity and (2) distance at peak velocity.

At larger aiming angles, longer distances were traversed at point of peak velocity (see Figure 5B), as shown by main effect of angle (BF_incl_ = 3.15 e+143). For peak velocity, whilst analyses suggested that peak velocity varied with aiming angles (main effect of angle: BF_incl_ =1.542 e+27), and that this effect depended on the session when the drug was ingested (Session x Drug x Angle interaction, BF_incl_ = 61.216), this was driven by somewhat larger peak velocities with straight ahead aiming (0°) than when aiming at different angles in the training session, predominantly for the Levodopa group (comparisons between 0° and all other aiming angles yielding cohen’s d values ranging between 0.233 to 0.414) than the placebo group (cohen’s d values ranging between 0.176 to 0.118).

**Figure 5.**
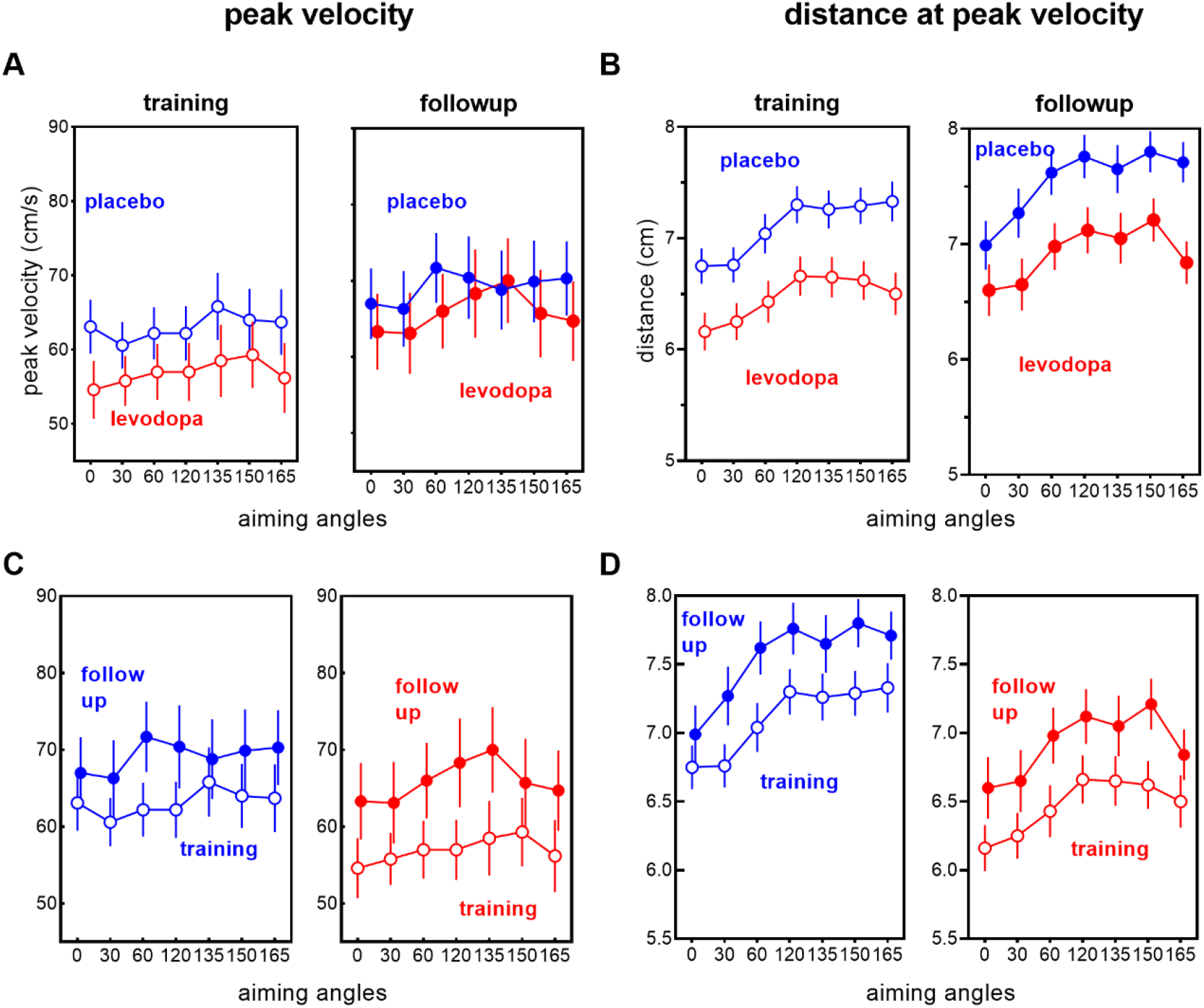
Levodopa reduced movement vigor, as shown by lower peak velocity (A) and lower distance at peak velocity (B) at training and follow-up. Vigor increased from training (clear circles) to follow-up (filled circles) Values are estimated marginal means and standard errors of the means.

In the training session where Levodopa was ingested, Levodopa reduced movement vigor, as shown by lower peak velocity (BF_10_ = 1.144 e+11) and reduced distance at peak velocity (BF_10_ = 2.209 e+57) see Figure 5). The finding that Levodopa decreased peak velocity and distance at peak velocity here appears to contradict previous findings demonstrating that levodopa increases movement vigor in effort-based decision-making tasks (Zenon et al., 2016), as well as in simple reaching tasks (Panigrahi et al., 2015). However, the decreased movement vigor with Levodopa shown here is consistent with recent work showing that optogenetic stimulation of midbrain dopamine neurons paradoxically decreased movement vigor during learning a skilled reaching task (Bova et al., 2020). Exogenous dopamine might thus have different effects on movement vigor with simple movements than with complex movements requiring motor skill (Bova et al., 2020), or when the task prioritizes movement accuracy, as in the current study.

Movement vigor also increased from the training session to the follow-up session (see bottom panels of Figure 5), as evidenced in a main effect of session for peak velocity (BF_incl_=1.58e+177), and distance at peak velocity (BF_incl_ = 8.37e+149).

## Discussion

Here, we explored the causal role of dopamine in motor learning in neurotypical individuals, by testing how the performance of goal-directed motor strategies was affected by dopamine during training, and by testing whether this manipulation altered performance at follow-up, when participants re-encountered the same task. We used a task with varying levels of difficulty, and incentivized accurate performance by making task success contingent upon accuracy. We found clear effects of exogenous dopamine increasing performance accuracy and lengthening deliberation time, particularly for difficult task conditions. Although the dopamine manipulation was only applied in the training session, the same pattern of behaviour (increased accuracy and deliberation time) was maintained at follow-up. Thus, in a skill learning task that incentivised accuracy, exogenous dopamine enhanced the tendency to trade speed for accuracy.

The finding that increasing dopamine availability causally increased accuracy in an explicit motor learning task that incentivised accuracy supports the view that dopamine influences decisions to engage instrumental motivation in motor learning. These results are consistent with reports that similar exogenous dopamine manipulations increase deliberative control (Wunderlich et al., 2012; Sharp et al., 2016). Indeed, the focus on explicit motor strategies here might be why we found effects of our dopamine manipulation, in contrast to previous null findings (Quattrocchi et al., 2018; Palidis et al., 2021). We provided participants with explicit instructions on how to aim by varying angles from a presented target, where larger aiming angles were associated with larger errors, lengthier reaction times, and were more difficult. In contrast, previous work rewarded participants for small, possibly implicit changes in behaviour (Palidis et al., 2021)), or used traditional paradigms that engaged mutually compensatory implicit and explicit motor learning operations (Quattrocchi et al., 2018)). Similarly, pharmacological manipulations of dopamine do not always result in observable effects on learning processes that have been characterised as reflexive or automatic (Wunderlich et al., 2012; Sharp et al., 2016; Grogan et al., 2017; Grogan et al., 2019). Indeed, studies which examine effects of dopamine manipulations on the relative contributions of deliberative, model-based learning and reflexive, model-free learning processes show selective effects of dopamine manipulations on model-based learning, and not model-free learning (Wunderlich et al., 2012; Sharp et al., 2016).

A second feature of this study that differed from previous work (Quattrocchi et al., 2018; Palidis et al., 2021) was the focus on manipulating task success. In contrast to previous work which manipulated reward by manipulating extrinsic rewards (points or money) contingent upon achieving some performance criteria (Galea et al., 2015; Steel et al., 2016; Quattrocchi et al., 2018; Spampinato et al., 2019), we focussed on success in achieving the task goal (target-hitting). Participants could only hit targets via accurate aiming, as the targets would move mid-movement by the required aiming angle. Task success thus required accurate performance. Although we provided auditory feedback for movements which met performance criteria, and this feedback could have been perceived as a form of extrinsic reward, Levodopa did not increase the number of movements that met the performance criteria. Our approach of making task success contingent upon accurate performance might have made our task more sensitive to the effects of exogenous dopamine. In contrast to direct manipulations of task success, which consistently yields clear effects on motor learning (Schaefer et al., 2012; Leow et al., 2018; Kim et al., 2019; Tsay et al., 2022), extrinsic rewards do not always alter motor learning (Steel et al., 2016; Spampinato et al., 2019) or motor performance (Grogan et al., 2020). Manipulations of task success might be more sensitive to effects of dopamine than manipulations of extrinsic rewards. Future studies could examine the role of dopamine in determining intrinsic motivation and sensitivity to extrinsic rewards by employing dopamine manipulations in behavioural paradigms which test the relative contributions of task success and extrinsic rewards (e.g., Vassiliadis et al., 2021).

Our finding that dopamine increases accuracy under more difficult task conditions is consistent with a view that dopamine guides instrumental motivation in *execution* of effort to attain accurate task performance. This extends previous research in humans, which has largely focussed on the role of dopamine in motivation to *select* effortful options, as the majority of previous studies have tested how dopamine manipulations alters decisions to select effortful options in tasks where each decision is only associated with a small or random probability of actually executing that response option (for a review, see Lopez-Gamundi et al., 2021). For example, in effort-based decision-making paradigms, methylphenidate, a dopamine and noradrenaline re-uptake inhibitor, increases young adults’ willingness to select more difficult working memory conditions (Westbrook et al., 2019). Dopamine denervation in Parkinson’s disease patients results in steeper discounting of effort for reward when choosing between effortful options in comparison to controls (Chong et al., 2015), and this deficit is partially remediated by medications which increase dopamine availability (McGuigan et al., 2019). By showing here that dopamine increased accuracy in a task where participants could not opt out of any trial, but could only choose how well to execute each trial, we infer that dopamine not only alters willingness to select effortful options, but also increases willingness to *execute* effort.

How might exogenous dopamine alter the engagement of instrumental motivation during skill acquisition? Influential theories suggest that slow “tonic” dopamine responses signal the background rate of reward, and drives motivation (Niv et al., 2007). Fast “phasic” dopamine responses signal unexpected outcomes and helps the actor learn to select choices that optimise outcomes (Steinberg et al., 2013). Recent studies however suggest that fast dopamine responses drive both motivation to work for rewards, and learning to select options leading to better outcomes (Hamid et al., 2016). Here, we used Levodopa, the precursor to dopamine, which increases dopamine availability within the brain. Although we do not fully understand how Levodopa affects fast and slow time-scale dopamine responses, animal studies have demonstrated that Levodopa increases phasic firing of striatal dopamine cells (Willuhn et al., 2014). We speculate that Levodopa employed at training increased accuracy by increasing the amplitude of phasic dopamine cell firing in response to target error feedback. Accuracy improvements that persisted at follow-up might have been associated with training-specific changes in excitatory synaptic transmission in the striatum that were altered by exogenous dopamine at training (Yin et al., 2009).

The increase in accurate aiming with Levodopa was accompanied by lengthier deliberation times, suggesting a more cautious mode of response overall – a classic *speed/accuracy tradeoff*. Intriguingly, increases in response caution have not been demonstrated in previous work examining how pharmacological manipulations of dopamine alters decision-making behaviour using evidence accumulation modelling (e.g., Winkel et al., 2012; Beste et al., 2018; Rawji et al., 2020). In those studies, dopamine manipulations either had no effect on response caution (Winkel et al., 2012), or increased response errors (Huang et al., 2015), or decreased response caution only in the context of proactive inhibition (Rawji et al., 2020). These mixed findings might be due to methodological differences, such as the use of different pharmacological agents (e.g., bromocriptine, a dopamine D2 agonist in Winkel et al., 2012; ropinirole, a dopamine D3 agonist in Rawji et al., 2020; and methylphenidate, a dopamine and noradrenaline re-uptake inhibitor in Beste et al., 2018) or the use of different tasks (e.g., simple decision-making tasks versus tasks requiring proactive inhibition in Rawji et al. (2020). Here, in a task that rewarded accuracy and not speed, exogenous dopamine resulted in participants trading-off speed for accuracy, particularly for difficult task conditions. Incentivizing both the speed and accuracy of movements can concurrently increase both speed and accuracy (Manohar et al., 2015). Future studies which manipulate dopamine whilst differentially incentivise speed versus accuracy during skill learning should elucidate how dopamine alters the trade-off between speed and accuracy in contexts where speed is prioritised (e.g., tennis) versus when accuracy is prioritised (e.g., laparoscopic surgery).

Skill learning is accompanied by a violation of the speed-accuracy trade-off, where both speed and accuracy are improved (Krakauer et al., 2019). Whilst both speed and accuracy of aiming movements improved at follow-up compared to training, we did not find evidence that dopamine modulated the size of the improvements in speed and accuracy. This might be due to the timing of the dopamine manipulation. Animal studies have shown that dopamine agonists and antagonists applied *before* learning have no effect on retention 24 hours later, but did alter retention when applied 3-6 hours *after* initial learning (Bernabeu et al., 1997). Similarly, dopamine medications ingested by Parkinson’s disease patients 8-24 hours *after* learning improve subsequent recall when tested a day after drug ingestion (Grogan et al., 2015). Dopamine might thus have a selective effect on offline consolidation of learning in a critical post-task period. Future studies examining how manipulating the timing of post-learning dopamine affects retention will elucidate the role of dopamine at different timescales in the motor memory consolidation.

Because we manipulated task success to incentivize accurate performance, we cannot make direct inferences about how dopamine alters “pure” intrinsic motivation to learn and perform motor skills. Increasing evidence demonstrates a key role of dopamine in learning in the absence of extrinsic rewards, and in self-evaluating performance without any extrinsic feedback (Ripollés et al., 2018; Duffy et al., 2022). For example, in zebra finches, dopamine neurons show spontaneous activity that correlates with song performance, even in the absence of extrinsic rewards, cues, or external perturbations (Duffy et al., 2022). Similarly, pharmacological manipulation of dopamine in humans can alter learning of novel words in the absence of explicit reward or feedback, and effects remain prominent on a no-drug follow-up session (Ripollés et al., 2018). To test the role of dopamine in intrinsic motivation in motor learning, future studies can combine pharmacological manipulations of dopamine with behavioural paradigms devoid of any form of extrinsic performance feedback, for example by removing or limiting visual feedback of movement (Vassiliadis et al., 2021), and/or having target appear only transiently at the start of the trial to preclude target-hits or target-misses (Taylor and Ivry, 2011).

Although we manipulated task success (i.e., target-hits in a target-hitting task), our paradigm does not allow us to dissociate if behaviour was motivated by avoidance of losses (target-misses) or pursuit of rewards (target-hits). A large body of knowledge demonstrates that rewards have potent effects in motivating effort (Takikawa et al., 2002; Xu-Wilson et al., 2009; Manohar et al., 2015; Manohar et al., 2017; Codol et al., 2019), even when rewards do not depend on how well we perform (Manohar et al., 2017), or when rewards are presented outside our conscious awareness (Pessiglione et al., 2008). Reward effects are evident even in very early visuomotor responses (Carroll et al., 2019b; Codol et al., 2023). Reward also makes us more willing to decide to engage costly cognitive resources, such as choosing to perform difficult working memory tasks, switch task sets, or inhibit prepotent responses (Westbrook and Frank, 2018). Whilst less is known about the motivational effects of punishment avoidance, it is clear that avoidance of negative outcomes is a potent driver of behaviour (Chen et al., 2020b). Whilst our evidence is broadly supportive of the proposal that dopamine alters motor performance by altering the brain’s sensitivity to rewards and increasing instrumental motivation, future studies should disambiguate if the effects shown here resulted from punishment avoidance or reward pursuit.

In conclusion, in a task which primarily engaged explicit learning processes, and incentivised accuracy by making task goals contingent upon accurate motor performance, we find that pharmacologically increasing dopamine availability increased performance accuracy, whilst concurrently increasing the amount of time used to deliberate and prepare movements. Effects were prominent both at training, and when re-encountering the same task after an overnight delay. We interpret the results as evidence supporting the view that dopamine plays a key role in decisions to engage instrumental motivation, which not only determines the quality of motor performance at initial learning, but also influences the quality of future motor performance when the same motor problem is reencountered. Whilst studies in animals and clinical populations have previously demonstrated a role for dopamine in motor learning (e.g.,Hosp et al., 2011; Isaias et al., 2011), this is, to the best of our knowledge, one of the first evidence for a direct link between dopamine and motor skill learning in neurotypical humans.

## Acknowledgements

The authors would like to thank Laura Fassbender for assistance with data collection.

## Notes

The authors report no conflicts of interest.

### Competing Interest Statement

The authors have declared no competing interest.

